# Densified Collagen Tubular Grafts for Human Tissue Replacement and Disease Modelling Applications

**DOI:** 10.1101/2021.02.05.429744

**Authors:** Alexander W. Justin, Federico Cammarata, Andrew A. Guy, Silas R. Estevez, Sebastian Burgess, Hongorzul Davaapil, Agavi Stavropoulou-Tatla, John Ong, Aishwarya G. Jacob, Kourosh Saeb-Parsy, Sanjay Sinha, Athina E. Markaki

## Abstract

There is a significant need across multiple indications for an *off-the-shelf* bioengineered tubular graft which fulfils the mechanical and biological requirements for implantation and function but does not necessarily require cells for manufacture or deployment. Herein, we present a tissue-like tubular construct using a cell-free, materials-based method of manufacture, utilizing densified collagen hydrogel. Our tubular grafts are seamless, mechanically strong, customizable in terms of lumen diameter and wall thickness, and display a uniform fibril density across the wall thickness and along the tube length. While the method enables acellular grafts to be generated rapidly, inexpensively, and to a wide range of specifications, the cell-compatible densification process also enables a high density of cells to be incorporated uniformly into the walls of the tubes, which we show can be maintained under perfusion culture. Additionally, the method enables tubes consisting of distinct cell domains with cellular configurations at the boundaries which may be useful for modelling aortic disease. Further, we demonstrate additional steps which allow for luminal surface patterning. These results highlight the universality of this approach and its potential for developing the next generation of bioengineered grafts.

## 1. INTRODUCTION

Bioengineered tubular grafts have the potential to address the growing clinical need for an effective and off-the-shelf source of replacement tissue. This notably includes blood vessel grafts (e.g. coronary and peripheral arteries, vascular access for hemodialysis) [1-5], but also grafts for the bile duct [6, 7], small and large intestine [8-10], oesophagus [11, 12], genitourinary [13, 14] and tracheobronchial [15-18] systems, and as guiding conduits for peripheral nerves [19]. Artificial tissue is also sought for disease modelling *in vitro* [20-22], such as to reproduce vascular pathology in atherosclerosis [23] and Marfan syndrome [24].

Tissue engineered grafts have been generated via processing of a wide range of synthetic polymers (e.g. polytetrafluoroethylene (PTFE), polyester, polyurethane), from cadaveric tissue, and via scaffold-free approaches [24-27]. Synthetic material-based grafts, while inexpensive and simple to manufacture, frequently lack the biological niche and degradation characteristics for cellular repopulation and remodelling. Furthermore, small diameter vascular grafts (< 6 mm) made from synthetic materials exhibit poor graft patency [28, 29]. Grafts derived from decellularized cadaveric tissue [30], while providing a high degree of bioactivity and innate vascularisation, are limited in availability and customisability in terms of geometry and composition. Further, complete removal of immunogenic foreign biomolecules remains a challenge despite advances in decellularization techniques [31]. To address some of these limitations, namely availability and customisability, an alternative approach is to seed cells upon a synthetic porous scaffold which subsequently produce native ECM to create human vessels [32, 33]. However, this method requires an expensive and lengthy production process, involving perfusion of large cell populations and subsequent decellularization. Scaffold-free approaches enable patient-specific graft generation and avoid the need for biomaterials which may impede the repair process [25]. However, they also require the culture of a dense population of cells for extended periods, and that the patient’s own cells be used, limiting their use as an off-the-shelf source of replacement tissue. Thus, generating implantable constructs using a cell-free method and employing fully definable and non-immunogenic biomaterials, remains a critically important challenge for the widespread use of bioengineered tubular grafts.

Despite the complications associated with using cells in graft generation, the manufacturing processes for producing more complex tissue types may need to be compatible with cell seeding both onto the surface and uniformly in the walls of the tissue, and may also require the incorporation of organoid structures to recapitulate tissue function, necessitating a benign manufacturing method. The grafts should nonetheless allow for cellular remodelling, demonstrate suitable and tuneable mechanical properties to match native tissue, and importantly be scalable from small animal models to human vessel dimensions. Additionally, tissue-specific challenges must also be addressed, such as thrombogenicity in blood vessels or acidity in urothelial conduits, and intestinal grafts will require fabrication methods to generate the high luminal surface area and rich vasculature for physiological function.

Bioengineered grafts have been produced via processing of biological polymers. Specific methods for producing collagen-based grafts include freeze-drying (yielding mechanically weak and highly porous sponges) [8, 34], electrospinning (high density of collagen, limited cellular remodelling and infiltration) [35], or through a hybrid with synthetic materials to provide the requisite strength for the collagen or ECM [36]. Alternatively, collagen hydrogel approaches support high bioactivity and consist of a macromolecular network of collagen fibrils which retain a significant volume of water; however, a standard collagen gel is mechanically weak and high collagen concentrations are difficult to deploy owing to high viscosity and rapid gelling times. Further, collagen fibrils are randomly orientated; in the native ECM, fibrils are hierarchically organised and aligned over distance which, alongside native crosslinking mechanisms, provide the tissue with mechanical support.

Several techniques [37-43] have been investigated to produce tubular collagen grafts by increasing the fibril alignment and density of a collagen hydrogel, thereby producing a mechanically strong construct while still retaining high bioactivity. This is achieved by removing a large fraction of the interstitial water from the hydrogel through *plastic compression* [38, 40], air drying [43], rotation-based water loss [42], gel compaction driven by cell traction forces [39, 44] or cyclic straining [37]. Tubular grafts are also typically produced by the rolling of densified collagen sheets (i.e. sutured into a tubular structure or rolled into spiral assemblies), but this leads to the formation of seams and variation of collagen density across the wall thickness [41, 45]. The resultant tubes tend to be thin-walled [37, 39, 40, 42], failing to match the wall thickness of human vessels, and are not easily scalable to human-sized vessels.

In this work, we describe a novel, materials-based method of manufacture for the formation of tubular grafts made of densified collagen hydrogel. The method allows for collagen densification by expelling interstitial water in both the axial and radial directions using a funnel-shaped mould. This results in mechanically strong tubes which are fully customisable in terms of tube diameter and wall thickness, yielding tubes for both small animal and human-sized tissues. Conduits with ultra-fine wall thicknesses can also be generated using pre-densified constructs and applying a secondary air-drying step. Tubes can be manufactured acellular, or cellularized by uniformly seeding cells into the walls of the conduit prior to collagen gelation and densification, or by seeding cells onto the luminal surface after the densification process is complete. To stabilise the collagen structure against acid degradation (e.g. urea) and to closer match the mechanical strength of real tissue, we further crosslink the densified collagen tubes using genipin, a naturally occurring and biocompatible crosslinking agent, which can be used in the presence of embedded cells. Alternative or additional steps to the standard technique enable more niche and disease modelling applications to be targeted. The shape of the funnel can be configured to produce a collagen tube with a collagen concentration gradient along its length. By sequential casting of cellularized collagen precursor solution, one can produce tubes with distinct cellular domains, which may be useful in disease modelling involving boundaries between cell types. Finally, a method has been developed for patterning the luminal surface which may support the future generation of intestinal crypt-villus structures.

## 2. RESULTS AND DISCUSSION

### 2.1. Rapid materials-based method of manufacture of densified collagen hydrogel tubes supports high customisability and off-the-shelf capability

We developed a novel method to transform low concentration collagen hydrogel, with randomly orientated fibrils, into an aligned, high concentration tubular collagen construct. This was achieved by casting collagen hydrogel into a funnel-shaped chamber (figure 1a,b) containing a central rod and withdrawing water from the gel via hydrophilic nylon membranes (1.2 µm pores) into wetted paper-towels at the bottom of the chamber, which led to the densification of collagen fibrils in the gel. The combination of a funnel-shaped chamber, water withdrawal through the base, and anchorage of the gel by the membranes, causes the solid phase of the collagen gel to contract into the lower cylindrical mould section. As the collagen gel contracts downwards, the reorganisation of the fibrillar structure leads to water being forced out of the pores, which collects on top of the densifying gel.

**Figure 1.**
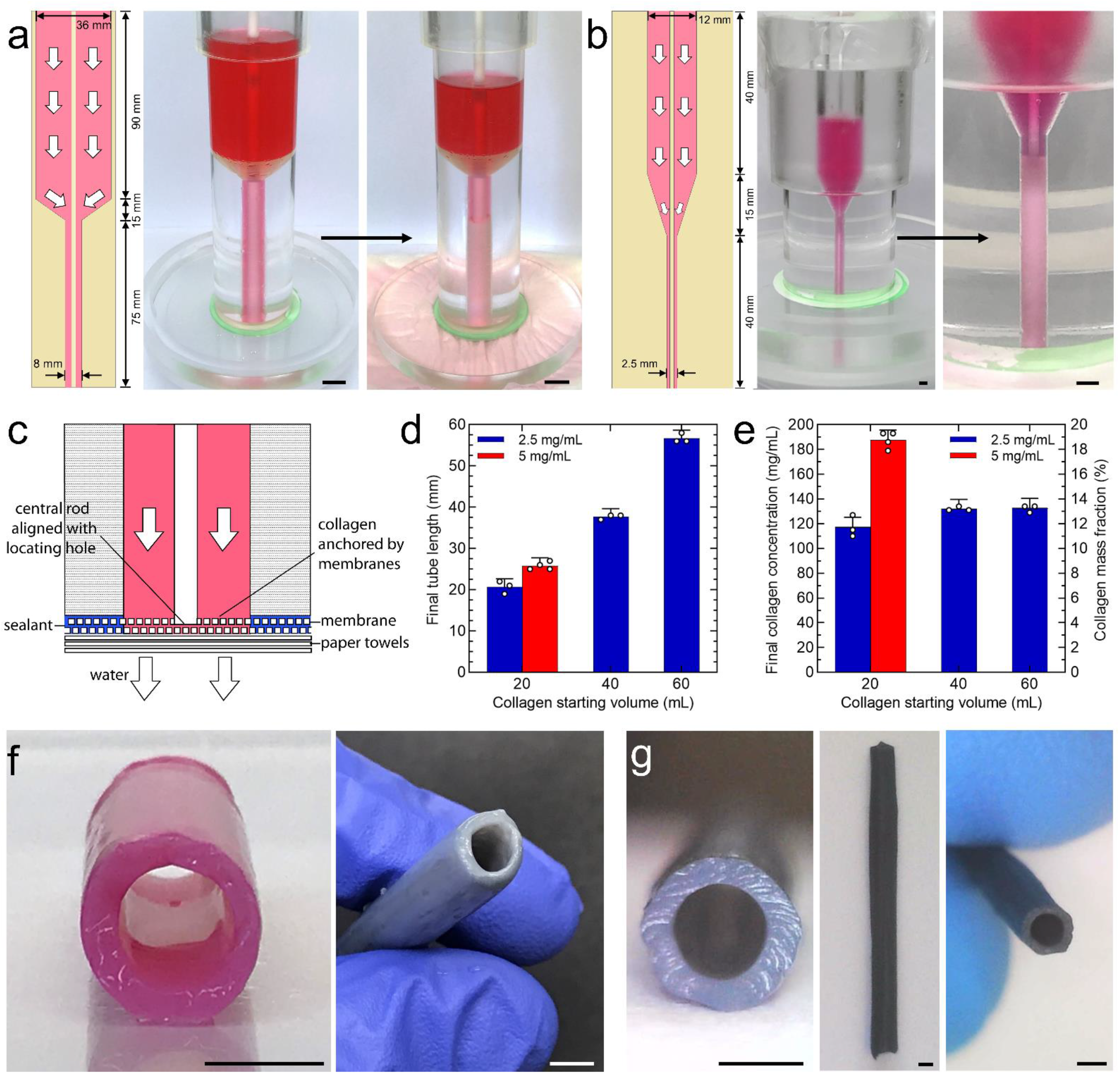
Densified collagen tube fabrication process. **(a)** Funnel used for generating densified collagen tubes with 5 mm luminal diameter and 1.5 mm wall thickness. The initial volume of collagen hydrogel was 60 mL. Scale bars: 8 mm. **(b)** Funnel used for generating densified collagen tubes with 1 mm luminal diameter and 0.75 mm wall thickness, prior to evaporative drying. The initial volume was 3 mL. Scale bars: 2 mm. **(c)** Schematic depiction of the funnel base used for production of densified collagen tubes, showing the bottom cylindrical section of the casting chamber, membrane configuration, central rod alignment, and water removal into paper towels. **(d)** Tube length as a function of collagen gel starting volume and concentration (5 mm lumen diameter and 1.5 mm wall thickness). Circles represent individual data points (*n* ≥ 3). **(e)** Final collagen concentration and mass fraction as a function of collagen gel starting volume and concentration (5 mm lumen diameter and 1.5 mm wall thickness). Final collagen concentration was calculated using final tube length (5 mm lumen diameter and 1.5 mm wall thickness). Circles represent individual data points (*n* ≥ 3). **(f)** Uncrosslinked (left) and genipin-crosslinked (right) densified collagen tube with a 5 mm luminal diameter and 1.5 mm wall thickness. Scale bars: 5 mm. **(g)** Genipin crosslinked densified collagen tube with 1 mm luminal diameter and ∼0.3 mm wall thickness, following air drying from 0.75 mm wall thickness. Scale bars: 1 mm.

The membrane assembly (figure 1c) both anchors the collagen gel at the base of the funnel-cylinder mould and impedes the rate of water removal, while also aligning the central rod in the chamber with a high degree of precision. This was necessary to produce tubes with fine (sub-millimetre) wall thicknesses. If no membranes are mounted at the bottom of the chamber, the collagen gel contracts rapidly upwards leaving an air pocket behind, which has the effect of preventing any further water removal from the hydrogel. Using membranes with significantly larger pore sizes leads to rapid water removal but causes the collagen gel to contract radially inwards, rather than downwards, preventing tube generation. Further, it is critical to the densification process that the collagen gel moves freely over the casting chamber surface, without significant friction or mass transfer of water outside of the gel surface; thus, a layer of oil lubricant is applied to the inside of the funnel. All water flow is therefore through pore percolation internal to the collagen gel.

Customizing the mould geometry enabled us to fabricate tubes of a wide range of different luminal diameters and wall thicknesses, and increasing the starting volume yielded proportionately longer tubes (figure 1d). By modulating the starting collagen concentration (e.g. 2.5 or 5 mg/mL), for a given starting volume, we produce tubes with different final collagen densities and mass fractions (figure 1e). The lower, 2.5 mg/mL collagen concentration may support faster cellular remodelling, while the higher, 5 mg/mL collagen concentration approaches the collagen content of native aorta (the collagen and ground substance is estimated in the literature at 47% [46]). Figure 1f shows a tube with a luminal diameter of 5 mm and wall thickness of 1.5 mm which was subsequently crosslinked with genipin, a naturally derived, water-soluble cross-linking agent. We also produced tubes of a significantly smaller luminal diameter (1 mm) and wall thickness (750 µm) which, through an additional evaporative drying step, could be processed to ∼300 µm wall thicknesses (figure 1g).

### 2.2. Densified collagen tubes exhibit strong collagen fibril alignment in the long-axis direction

The tubular wall of the densified collagen tubes, in both hydrated and dried states, was characterized using second harmonic generation (SHG) imaging and scanning-electron microscopy (SEM) respectively (figure 2a,b). These imaging techniques revealed a network of interconnected fibrils and fibres (bundles of densely packed fibrils), predominantly aligned in the axial direction (i.e. the long axis of the tubes). The 2D fibril angle orientations obtained from the SEM micrographs are illustrated in figure 2c for 1 and 10 mM crosslinked collagen tubes, with 2.5 mg/mL starting collagen concentration (for a description of the method employed see supporting information *SEM image analysis* and figure S1). Higher magnification SEM images such as figure 2d show regions of fibril bundles, which are separated and connected by regions of more sparse bundles and individual collagen fibrils. Estimation of the fibril diameter using such images showed a modal average in fibril diameter of 50 and 65 nm for 1 and 10 mM crosslinked collagen tubes, respectively (figure 2e and supporting figure S2). In this orientation, the collagen networks have such a high pore connectivity that it is difficult to delineate any individual pores. When sectioning across the tube wall (figure 2f), the collagen fibril density was found uniform across the thickness of the wall. The pores in cross-section were better defined; low magnification SEM images (figure 2g) were used to obtain pore area distributions (figure 2h), showing a modal average pore area of 0.2-0.4 µm^2^ (supporting figure S3). It should be noted that the fibril diameter and pore spaces were estimated using dried collagen tube sections rather than the standard hydrated collagen gel tubes, so deviations are expected. However, the drying method employed kept ice crystal formation to a minimum (see Methods – Scanning electron microscopy).

**Figure 2.**
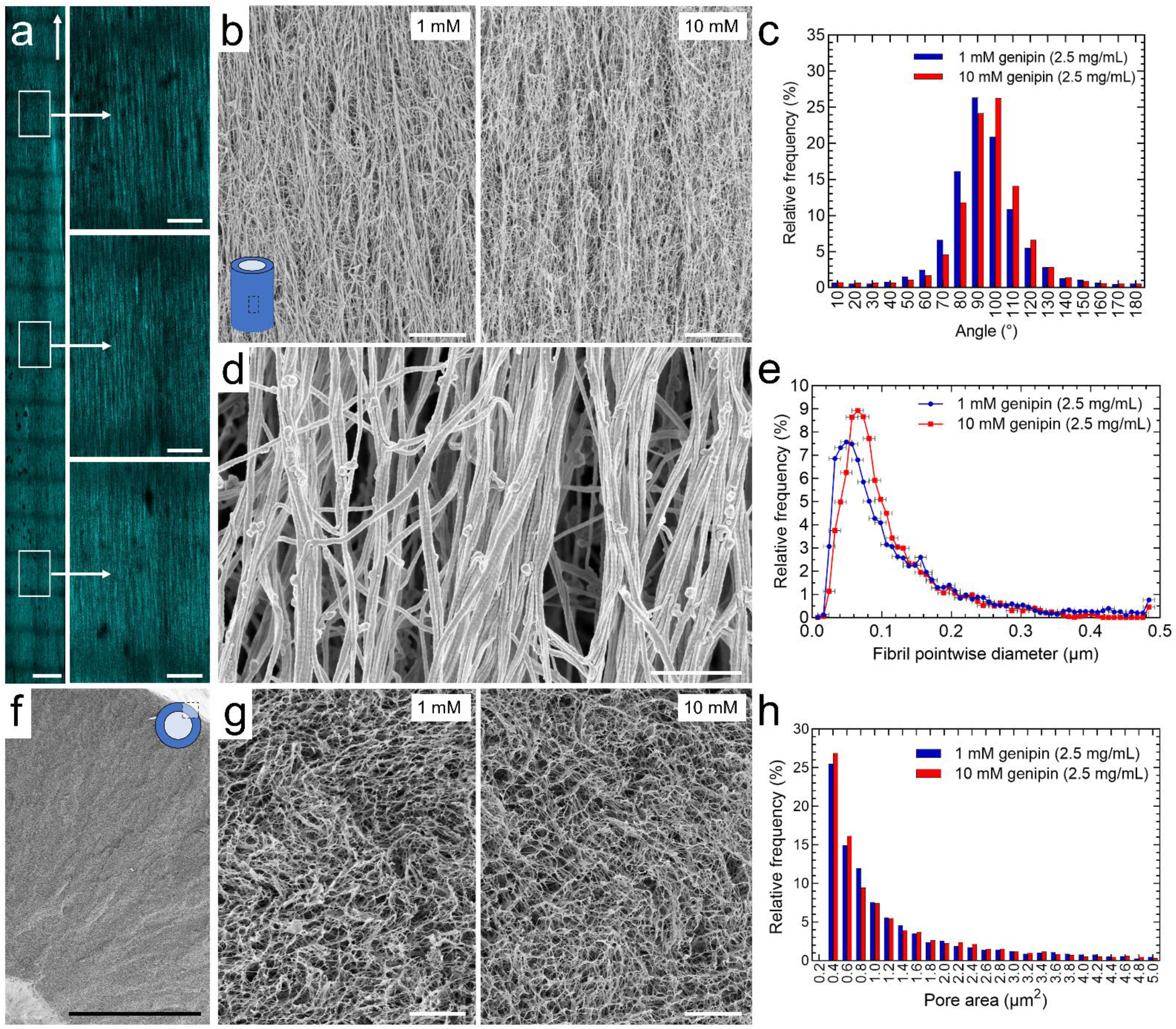
Densified collagen tubes exhibit strong fibril alignment. **(a)** Representative image of uncrosslinked densified collagen tube in the wet state displaying fibre orientation and uniform fibre density, as shown using second harmonic generation microscopy. Arrow denotes long axis of the tube and insets show higher magnification images. Scale bars: 500 µm (low magnification image); 100 µm (high magnification images). **(b)** Low magnification SEM micrographs of the freeze-dried outer surface of densified collagen tubes, crosslinked using 1 and 10 mM genipin solutions. Scale bars: 10 µm. **(c)** Distributions of 2D fibril orientation angles (with respect to the horizontal) for 1 and 10 mM crosslinked densified collagen tubes using data obtained from the low magnification SEM images shown in b (histogram bins right closed). Curved fibrils are sampled at every point along their length, so they contribute to each orientation by the fraction of their length that is at that orientation. **(d)** Typical high magnification SEM micrograph of the outer surface of 1 mM densified collagen tube. Scale bar: 1 µm. **(e)** Distribution of fibril diameters using data obtained from high magnification SEM images such as the one shown in d (histogram bins right closed). **f and g)** Low and high magnification SEM micrographs of the tube wall in cross-section. Scale bars: 500 µm (low magnification); 10 µm (high magnification). **(h)** Distribution of pore areas using data obtained from high magnification SEM images such as those shown in g (histogram bins right closed). SHG imaging and SEM were undertaken using tubes with 2.5 mg/mL starting collagen concentration.

This densification process enables graft production with collagen content similar to the high collagen density of native tissues (e.g. aorta [46]), and the collagen fibril density was constant across the wall thickness and along the tube length. The resulting tubular constructs display a significant alignment of collagen fibrils parallel to the long axis of the tubes and the hierarchical structure (individual fibrils and fibril bundles (i.e. fibres)) found in native tissues. In blood vessels, the collagen fibrils are orientated predominantly in the hoop (circumferential) direction to accommodate pulsatile flow. It may be possible to influence the collagen fibril alignment in our tubes by tailoring the funnel geometry and by reducing individual fibril lengths, while improving bulk cellular migration by modulating the fibril bundling process[47] in order to increase the average pore size. Furthermore, we postulate that once implanted, the collagen construct will be remodelled over time through cellular degradation processes and ECM deposition to better imitate the intricate structure of the native vessel, all while maintaining functionality as a viable tubular graft.

### 2.3. Genipin-crosslinked densified collagen tubes exhibit functional properties suitable for implantation

While the densified collagen tubes display suitable mechanical strength for handling and cell seeding, we used a genipin crosslinking step to better imitate the mechanical strength of real tissue and eliminate any swelling and dissolution behaviour of collagen in acidic conditions (i.e. for applications where the perfusate is acidic). The genipin crosslinking was performed at a biocompatible concentration of 1 mM, suitable in tubes where cells have been pre-seeded into the walls of the scaffold, or 10 mM, producing mechanically strong structures for burst pressure and suture retention testing.

We measured the axial strength of uncrosslinked and crosslinked tubes under quasi-static uniaxial tension using samples extracted from 2.5 and 5 mg/mL collagen tubes (5 mm luminal diameter, 1.5 mm wall thickness). Figure 3a shows that the tube strength increases with starting collagen concentration and crosslinker concentration. The peak forces during suture pull-out were measured for 10 mM genipin-crosslinked tubes, with 2.5 and 5 mg/mL starting collagen concentrations, which were normalised by the tube wall thickness (figure 3b). We also measured the maximum pressures at bursting point of these tubes (figure 3c). Figure 3c also shows that the pressurised tubes fail due to a crack propagating in the axial direction, normal to the largest (hoop) stress. As expected, the 2.5 mg/mL tubes exhibit lower peak forces during suture pull-out and lower burst pressures, when compared to the 5 mg/mL tubes.

**Figure 3.**
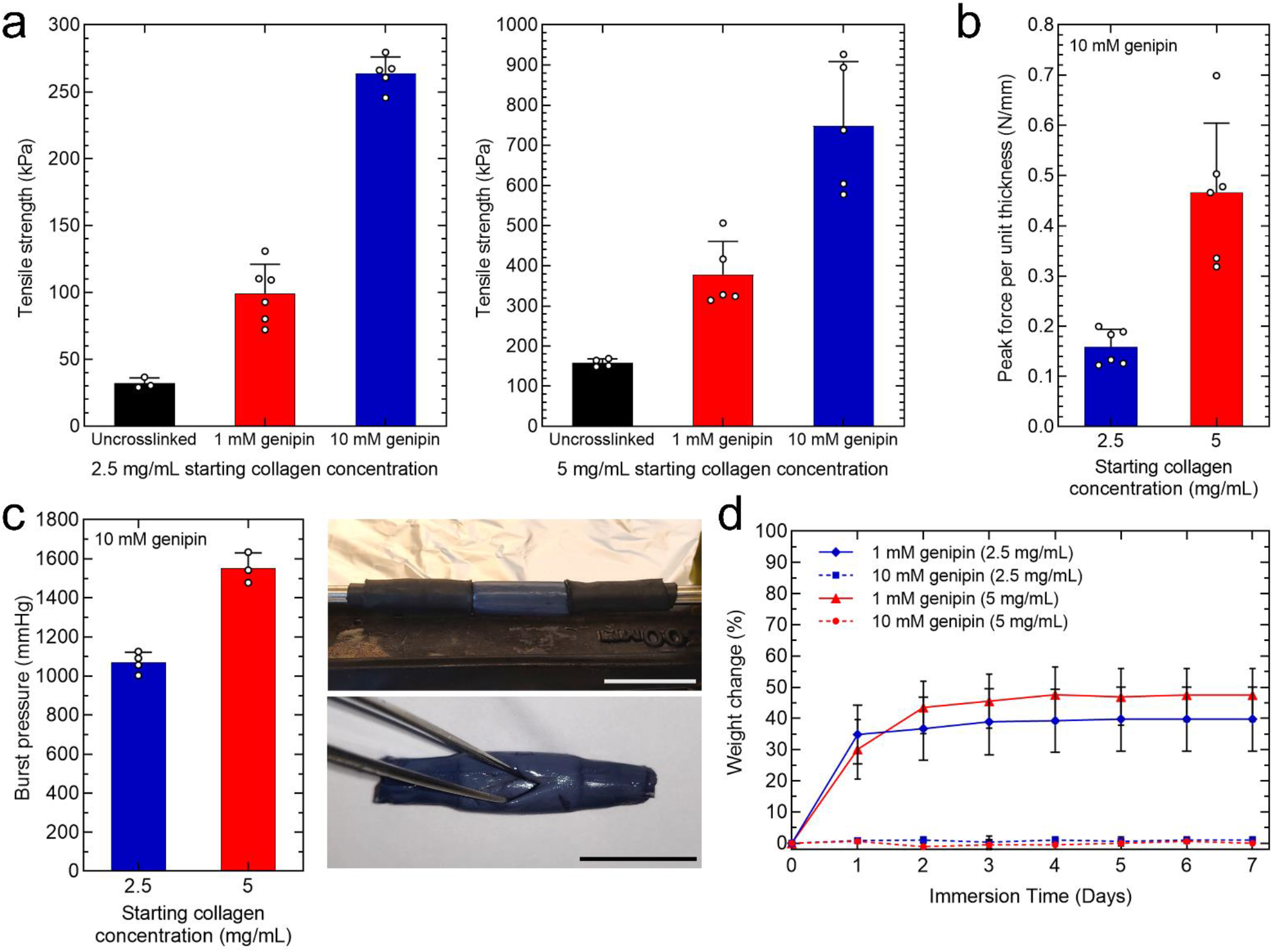
Genipin crosslinked collagen tubes exhibit useful properties for implantation. **(a)** Axial tensile testing of dogbone specimens (*n* ≥ 3) extracted from the walls of densified collagen tubes with 2.5 and 5 mg/mL starting collagen concentrations. **(b)** Peak forces during suture pull-out normalised by the tube wall thickness for 10 mM genipin-crosslinked tubes (*n* = 6). **(c)** Burst pressures for 10 mM genipin-crosslinked tubes (*n* ≥ 3). Images show heat shrink tubing used to seal the ends onto stainless steel hypodermic tubing (top), and a burst tube (bottom) due to a crack propagating in the axial direction. Scale bars: 20 mm. **(d)** Acid solubility data obtained from immersion in 100 mM acetic acid solution showing percentage change in weight over a 7-day period for 1 and 10 mM crosslinked tube samples (*n* ≥ 3). 5 mm diameter sections were extracted using a biopsy punch from tubes of 5 mm luminal diameter and 1.5 mm wall thickness. Bars (a-c) and data points (d) show arithmetic mean ± standard deviation; circles (a-c) represent individual data points.

Crosslinking with genipin prevented collagen dissolution and reduced collagen swelling (figure 3d). 1 mM genipin-crosslinked tubes, with 2.5 and 5 mg/mL starting collagen concentrations, underwent swelling in the first 1-2 days when immersed in a 100 mM acetic acid solution, after which a plateau was reached. Crosslinking with 10 mM genipin eliminated the swelling of the tubes entirely.

Direct comparison with previous work is hampered by a number of disparities, most notably in the testing configuration (e.g. tube, sheet, spiral assemblies) and loading conditions (e.g. loading rate, a priori cyclic straining to further strengthen the collagen, wet or quasi-dry state). Suture pullout data are reported in terms of force, so the values are influenced by the suture size and specimen geometry (e.g. for a given suture gauge, a thicker sample is expected to result in a larger peak force during pullout). Although our tube properties, namely axial tensile strength and burst pressures, are inferior to those reported for native blood vessels (1.5-3 MPa [48-50] and 1600-2500 mmHg [51, 52]), the tubes approach mechanical properties suitable for implantation; tensile strength values are of the order of 1 Mpa and burst pressures in the range of 1500 mmHg (5 mm luminal diameter, 1.5 mm wall thickness). For thin-walled cylinders, the relationship between internal pressure and the stresses in the wall can be readily derived by means of a force balance. The pressure is inversely proportional to the tube diameter, hence smaller diameter tubes would exhibit higher burst pressures for a given wall thickness and hoop stress. Additionally, fibril alignment in the hoop direction would increase the hoop stress and hence the burst pressure. Regarding suturability of our tubes, it has been previously demonstrated [6] that mouse-sized densified collagen tubes, when used as a replacement for the common bile duct, have sufficient mechanical strength for surgical implantation and function.

Future work will investigate the incorporation of other biopolymers, such as elastin, into the densified collagen structure. Elastin enables arteries to elastically expand and contract to accommodate pulsatile flow; however the addition of soluble elastin, added to the precursor collagen gel solution, was found to have a negligible effect on the tubes, even when genipin crosslinking was applied. It may be possible to use higher molecular weight elastin in the future which imparts a greater influence upon the densified fibril structure.

### 2.4. Manufacturing method supports luminal surface cell seeding and uniform cell seeding into tube walls

We seeded fluorescent hepatic epithelial cells (HepG2s) onto the luminal surface of the collagen tubes which, under static culture conditions, produced a monolayer structure after 1 week (figure 4a). In addition, we embedded cells into the walls of the densified collagen tubes by mixing them into the collagen gel precursor solution (figure 4b). When the collagen was gelled and densified, this led to a uniform cell distribution within the tube walls, as demonstrated by experiments using populations of rat aortic smooth muscle cells (SMCs) and HEPG2s (figure 4c).

**Figure 4.**
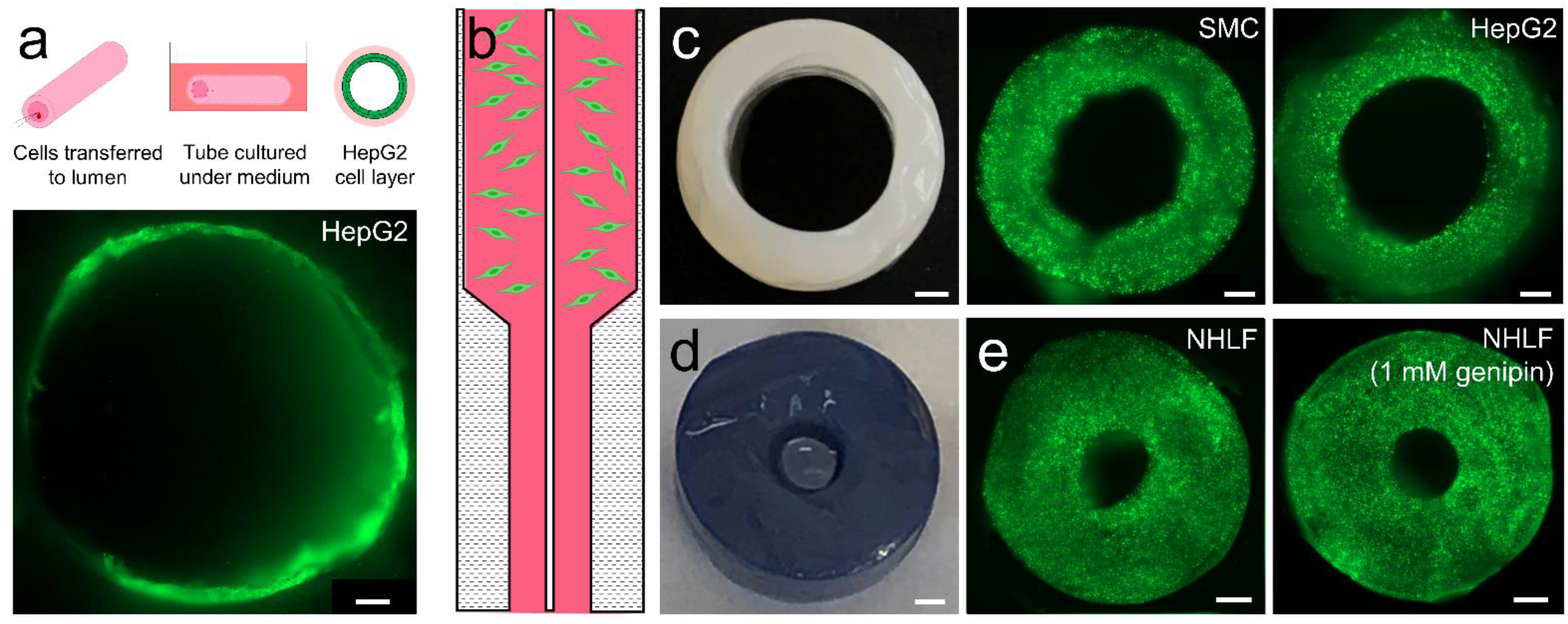
Densified collagen tubes support cell seeding onto luminal surface and into tube walls. **(a)** Schematic of surface seeding process. HepG2s were pipetted into lumen of the tube (5 mm luminal diameter, 1.5 mm wall thickness) and statically cultured for 1 week. This led to formation of a cell monolayer on the luminal surface. Scale bar: 200 µm. **(b)** Schematic depiction of the bulk cell seeding process. Cells were mixed into the upper part of the gel precursor solution. **(c)** Cross-sectional image of 5 mm luminal diameter, 1.5 mm wall thickness, uncrosslinked tube. SMCs and HepG2s were encapsulated into the tube walls. Scale bars: 1 mm. **(d)** Densified collagen tubes after crosslinking exhibit characteristic dark blue colour. Scale bar: 1 mm. **(e)** Samples of NHLF-seeded collagen tubes, with 2 mm luminal diameter and 3 mm wall thickness, before and after 1 mM genipin crosslinking (24 h), showing high viability of cell population. All cells were stained using calcein AM (green). Scale bars: 1 mm.

Genipin crosslinker solutions caused a significant colour change in the densified collagen tubes (figure 4d). To test short-term cell response to genipin, we immersed a cellularized densified collagen tube with a large (3 mm) wall thickness (figure 4e) in cell medium containing 1 mM genipin for 24 h and showed that populations of normal human lung fibroblasts (NHLFs) maintained their viability over this time period, under static culture conditions.

For surface cell viability, SMCs were seeded onto 1 and 10 mM collagen tube substrates, alongside tissue culture plastic (TCP) and polydimethylsiloxane (PDMS) which served as positive and negative control surfaces respectively. Figure 5a shows the percentage of alamarBlue reduction on the surface-seeded substrates, measured on alternate days, from Day 1 to Day 13. This reduction increased with culture time on the 1 and 10 mM crosslinked collagen tubes suggesting that the SMCs maintained a reducing environment within their cytoplasm. We observed a negligible difference in the alamarBlue reduction between the 1 and 10 mM crosslinked collagen tubes, and the TCP control. Cells reached confluency at Day 11.

**Figure 5.**
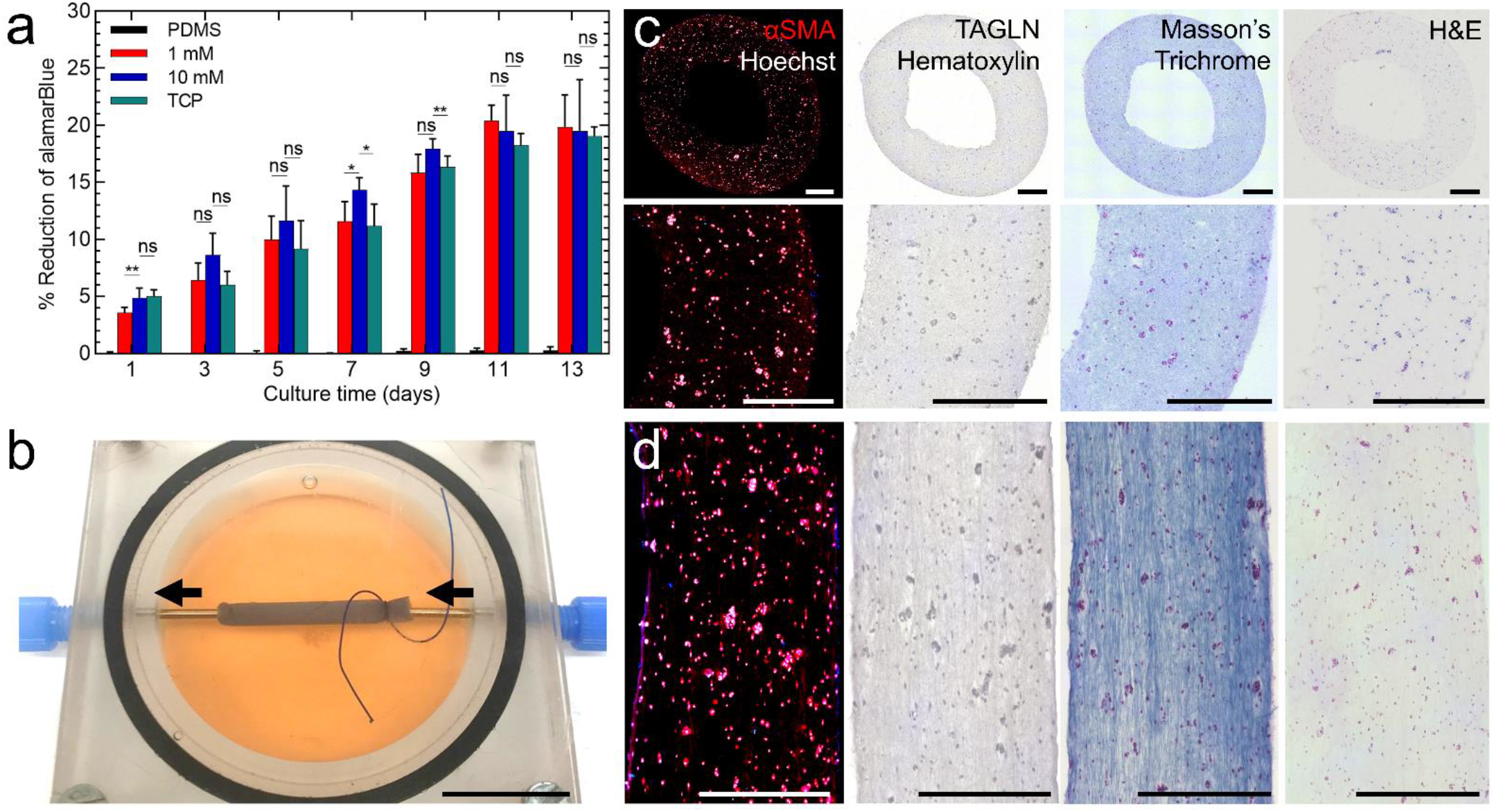
Cellularized densified collagen tubes under static and perfusion culture. **(a)** Surface cell viability was measured up to day 13 using an alamarBlue assay. Negligible statistical difference was observed between 1 and 10 mM crosslinked-samples, and between the tube samples and the tissue culture plastic (positive control). Statistical significance was analysed using the Kruskal-Wallis and the Games-Howell *post hoc* test. Data are presented as the arithmetic mean ± standard deviation; *n* = 4; \**p* < 0.05, \*\**p* < 0.01; ns means no statistically significant difference. **(b)** Closed loop perfusion of densified collagen tubes with cells embedded in the walls. Flow rate was maintained at 600 mL/hr for 10 days using a peristaltic pump. Scale bar: 20 mm. **(c)** Immunohistochemical and histological staining of tube cross-sections at Day 10. α-SMA expressed by SMCs in red, Hoechst counterstain in white, alongside TAGLN expression by SMCs with Hematoxylin counterstain. Evaluation of ECM and embedded cells by Masson’s Trichrome (collagen in blue, cells in purple, cell nuclei in dark blue); and H&E staining (ECM and cytoplasm in pink, cell nuclei in purple). Cross-sections display a high density of uniformly distributed cells. Scale bars: 500 µm. **(d)** Similar staining of sections of cellularized densified collagen tubes in the long axis direction, showing some cell elongation. Scale bars: 500 µm.

### 2.5. *In vitro* perfusion study demonstrates long-term survival of a high density cell population in the walls of densified collagen tubes

High densities of cells embedded in thick-wall constructs necessitate a perfusion culture system for maintenance of cell viability. Densified collagen tubes, containing 5 million SMCs at a cell density of 25 million SMCs/mL and crosslinked with a 1 mM genipin solution, were perfused at 600 mL/hr for 10 days using a custom-made perfusion chamber (figure 5b) to demonstrate cell viability. Immunohistochemical and histological staining of tube sections (figure 5c,d) demonstrated both the survival of a high density of cells within the walls of a tube under perfusion culture conditions and showed the proliferation of these cells within dense collagen. Through representative cell counting of fixed 5 µm cross-sectional slices, we measured a total cell number of 12 million SMCs, yielding a final cell density of 250 million SMCs/mL. Staining cross-sectionally (figure 5c) and in the longitudinal axis direction (figure 5d) showed no variation in cell density across or along the collagen walls, further demonstrating uniform cell seeding. We also detected some limited elongation of the SMCs in the axial direction which, given the alignment of the collagen fibres and anisotropy of the pores, is to be expected. Alongside cell uniformity, the Masson’s trichome stain demonstrated the alignment of collagen fibrils in the longitudinal direction and showed a uniformity in collagen density across the tube wall.

Since the densification process reduces the gel volume significantly (by a factor of 20-50, dependent on the funnel geometry and starting collagen concentration), the cell density can also be substantially increased which has the potential in future to reach physiological levels. While in this work we perfused crosslinked tubes, it is also possible to delay genipin crosslinking since the tubes can mechanically support themselves; this may be desirable as an uncrosslinked tube can undergo faster cellular remodelling in the short-term. Biocompatible genipin crosslinking can subsequently be performed at any stage thereafter.

### 2.6. Densified collagen tubes with multiple cell domains exhibit a non-trivial boundary between cell populations

Disease modelling applications may require the formation of a densified collagen tube with two distinct regions consisting of two different cell types (e.g. two lineage-specific types of SMC). For these tubes, we used a mould with sloped walls (figure 6a) and postulate that the funnel shape is important to generating different boundaries between the cell populations. Thus, we manufactured tubes in which we cast half the precursor solution with one cell type and cast the second half of the precursor solution with the second cell type (figure 6b). Cells from each sub-population were maintained in place by the relatively high viscosity of the precursor gel solution prior to gelation. Imaging revealed two distinct cell domains consisting of either green or red fluorescently labelled HUVECs (figure 6c). Interestingly, the non-trivial mapping of the initial undensified collagen gel in the funnel section to the final densified collagen tube in the cylindrical section was such that the top group of HUVECs was located around the lumen while the bottom group of HUVECs populated the outer part of the walls, near the boundary between the cell domains (figure 6d). The exact configuration of the cells in the final tube is likely determined by the exact angle of the sloped funnel and could be investigated further in the future. Second harmonic imaging of the tubes formed using the sloped funnel showed density variation along the tube length (supporting figure S4).

**Figure 6.**
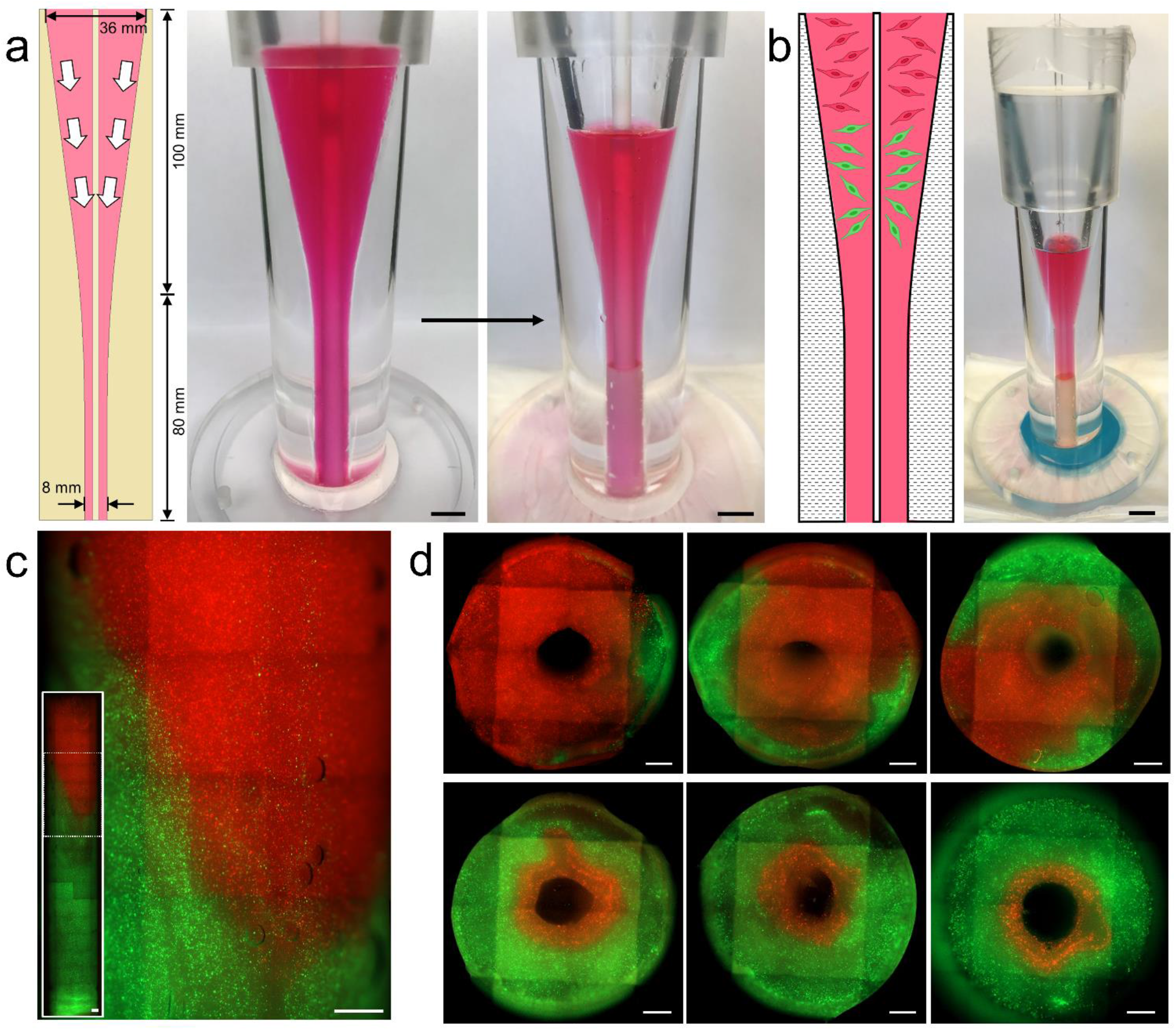
Sloped funnel densification for disease modelling applications. **(a)** Sloped funnel densification process for disease modelling applications. Images are taken before and after collagen densification. Scale bars: 8 mm. **(b)** Schematic showing position of green and red fluorescently-labelled cells (GFP and RFP HUVECs respectively) in undensified collagen gel, alongside image of densified collagen tube containing fluorescently-labelled cells. Scale bar: 8 mm. **(c)** Images showing the interface between two cell domains in densified collagen tube in the longitudinal axis direction. Inset shows whole collagen tube. Scale bars: 1 mm. **(d)** Cross-sectional views moving down the densified collagen tube, showing transition from RFP to GFP HUVECs. Note that due to the densification process, RFP HUVECs are mapped from the top of the gel to wrap around the lumen, even in the GFP HUVEC region of the tube. Scale bars: 1 mm.

While a density gradient may not be useful for implantable tissue equivalent structures, the multi-domain tubes which make use of the slope-walled funnel moulds may find future application as a means of studying aortic aneurysms. The mapping of volume elements of collagen gel from the funnel into the final cylindrical form is non-trivial; elements near the central rod are drawn down inside the surrounding gel due to the shape of the funnel. This results in the observed spatial organisation of the domains near the boundary, with one cell type populating the region near the lumen and the other type populating the outer wall region, which is highly reminiscent of the pattern of distribution of SMCs in the proximal aorta at the boundary between SMCs of neural crest and lateral plate mesoderm origins [53]. Indeed, this overlapping pattern is thought to be one of the reasons why aortic aneurysms and dissection are prevalent in the aortic root and ascending aorta [54]. Therefore, this manufacturing technique may in future allow us to recreate the distinct cellular spatial configurations in the proximal aorta, notably by varying the funnel geometry, and heterotypic SMCs could be seeded in distinct overlapping domains in our grafts to investigate if proximity of different SMC lineages is detrimental to graft integrity [55].

### 2.7. Luminal surface patterning of densified collagen tubes produces structures reminiscent of intestinal villi

Intestinal applications of the tubes may require the generation of crypt-villus structures on the luminal surface to promote efficient exchange of nutrients. To this end, we developed a process for patterning 3D printed structures in gelatin hydrogel (figure 7a-c) which, following casting and densification of the collagen, generated a regular pattern of villi-like structures on the luminal surface, with widths of 250 µm and heights of 500 µm (figure 7d,e). However, large extrusions from the surface can hold up the densification process as the collagen gel is unable to deform around such features.

**Figure 7.**
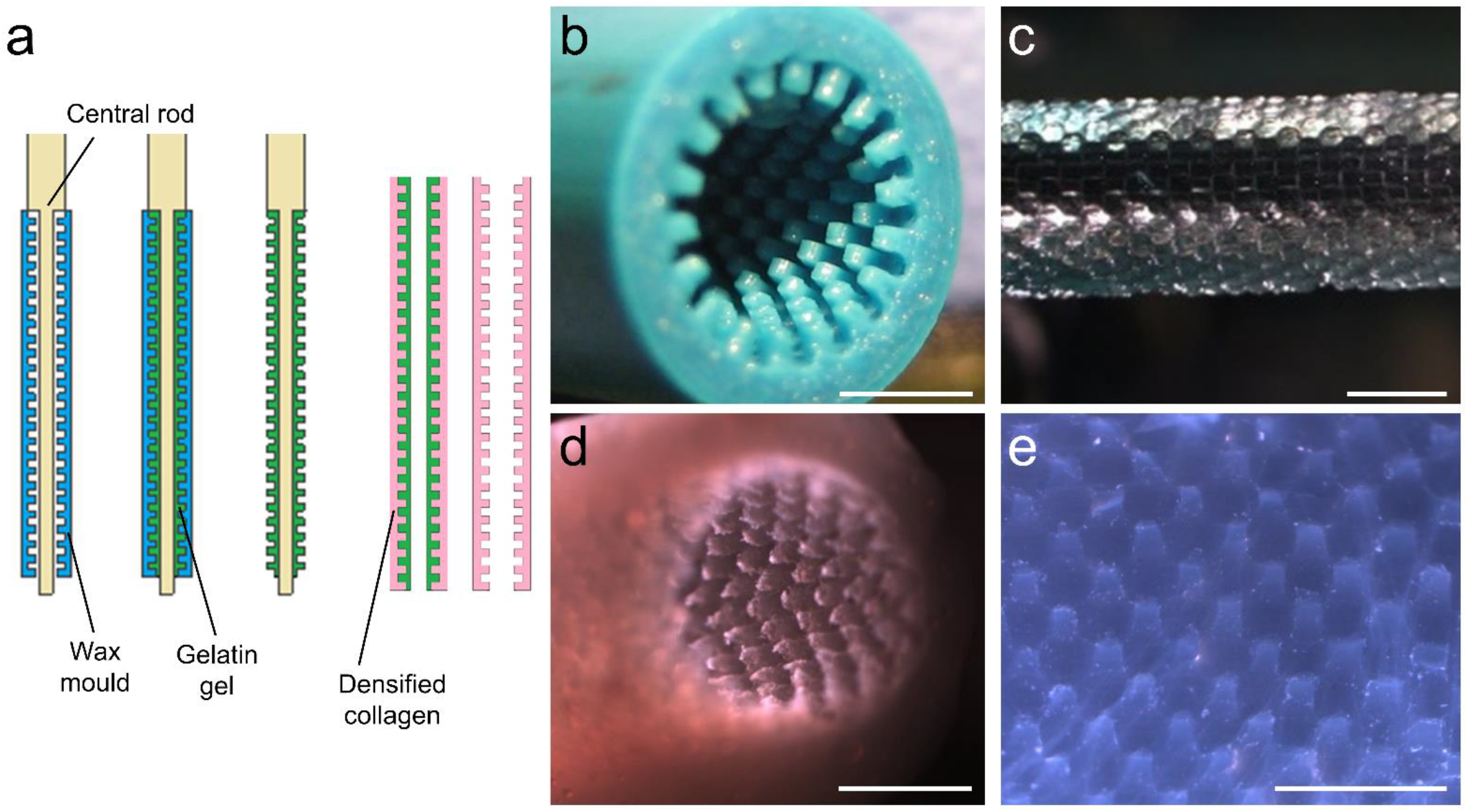
Luminal surface patterning of densified collagen tubes. **(a)** Schematic showing luminal surface patterning process. Wax mould (blue) is mounted on a central rod with a cutaway. Wax mould is filled with gelatin gel (green) and the wax mould removed. After densification, the central rod is removed, and the gelatin melted out of the densified collagen tube (pink). **(b)** 3D printed wax mould with patterned luminal features. Scale bar: 2 mm. **(c)** Gelatin gel inverse features, mounted on central rod. Scale bar: 2 mm. **(d)** Densified collagen tube with luminal surface features. Scale bar: 2 mm. **(e)** Luminal surface features at higher magnification. Scale bar: 2 mm.

## 3. CONCLUSIONS

We present a materials-based method of manufacture for generating bioengineered tubular constructs from densified collagen hydrogel, through an approach which does not require the use of cells or decellularization methods, supporting true *off the shelf* capability. The grafts are highly customizable in terms of diameter and wall thickness, seamless, and display a uniform fibril density across the wall thickness and along the tube length. Through genipin crosslinking, we show that we can eliminate collagen swelling and dissolution in acidic conditions, and generate constructs with the requisite mechanical strength for implantation. Critically, our method enables rapid and inexpensive fabrication (i.e. a few days), as opposed to methods relying on cell seeding, matrix deposition, or bioreactor culture. While acellular grafts can be readily produced, the cell-compatible densification method also enables a high density of cells to be uniformly seeded into the tube walls during fabrication, as well as onto the luminal surface. In addition, by varying the mould geometry, we can produce tubes with a collagen density gradient which, when coupled with distinct cell domains, may be useful in studying aortic diseases. We therefore conclude that our method represents a promising strategy for the generation of conduits for tissue replacement and disease modelling applications. Future work will focus on an assessment of our grafts as an interposition graft in small and large animal models.

## 4. EXPERIMENTAL SECTION

### Materials

Collagen hydrogel was generated using an established method [56] using rat tail tendon collagen solution (10 mg/mL, Corning), dissolved in 0.02 N acetic acid. Briefly, for preparation of 2.5 mg/mL collagen precursor solution (20 mL), the reagents used were: 10x M199 medium (2 mL, Sigma), 1 M NaOH (0.250 mL), stock 10 mg/mL collagen solution (10 mL), sodium bicarbonate solution (0.6 mL), penicillin-streptomycin (0.2 mL, Sigma), amphotericin-B (0.2 mL, Sigma), and cell medium (6.75 mL). Viscous collagen solution was transferred using a syringe and added last. The solution was shaken vigorously until thoroughly mixed and centrifuged at 2000 g for 5 min at 4 °C.

Perspex chambers, for casting and collagen densification, were machined to tube specifications, consisting of a funnel and a flange base. Additionally, top and bottom jig pieces were machined from Perspex, and PTFE and styrene rods were purchased to size (Engineering & Design Plastics). Nylon membranes were used with pore sizes of 1.2 µm (Millipore).

### Fabrication method of densified collagen tubes

Funnels and additional components were immersed in 70 % ethanol and dried prior to experiments. Internal funnel surfaces were coated in sunflower oil and positioned upside down. Using a biopsy punch (Agar Scientific), a hole was made in a 1.2 µm nylon membrane to match the diameter of the central rod. Using a machined jig, the locating hole in the membrane was positioned centrally over the end of the funnel and mounted using dental silicone (Dental Sky). An additional nylon membrane was then mounted to produce a two-layer membrane configuration. Membranes and funnel were subsequently pre-wetted with sterile phosphate-buffered saline (PBS, Sigma) to liberate any trapped air bubbles between the membranes. A water-impermeable polystyrene sheet was placed under the membranes to prevent early leakage of the collagen precursor solution and was held in place with the jig.

Collagen precursor solution was prepared as described above and transferred to the funnel. The collagen solution was visibly clear of air bubbles after centrifugation; failure to remove all the bubbles led to sub-optimal tubes with high numbers of defects or could disrupt the densification process entirely. The collagen solution was poured very slowly down one side of the internal funnel surface, thereby preventing an emulsion of oil droplets in the collagen solution and preventing any turbulence which may trap air bubbles. The pre-wetted membranes allow the infiltration of collagen solution, which subsequently provided an anchoring surface for the collagen gel. Once the whole collagen solution was in the funnel, the central rod, also coated with sunflower oil, was slowly lowered into the chamber, and interlocked into the locating hole in the top membrane, providing alignment to the rod from below. A second Perspex jig provided central alignment of the rod from above.

The whole assembly was subsequently left in a 37 °C incubator for 1 h to allow the collagen to gel. Following this, the bottom jig was removed along with the water-impermeable polystyrene sheet, and the funnel assembly positioned on top of pre-wetted paper towels. The full assembly was then placed in a 37 °C incubator, or under a biological cabinet at room temperature, for 1-24 h for the collagen gel to densify (depending on collagen volume and funnel geometry). After densification, the densified collagen tube was carefully extracted from the lower cylindrical section of the funnel. To undertake the optional evaporative densification step, the rod-tube assembly was suspended horizontally and rotated under a biological safety cabinet for 3-4 h.

### Crosslinking

Genipin (Bioserv UK) was resuspended in sterile PBS and prepared at 1 and 10 mM concentrations. Densified collagen tubes were immersed in genipin solutions and placed on an orbital shaker for 24 h at room temperature, followed by a further 24 h in sterile PBS. Collagen tubes containing embedded cells were crosslinked by immersing the constructs in cell medium containing 1 mM genipin, and incubating at 37 °C for 24 h.

### Tube characterisation

#### Scanning electron microscopy

Densified collagen tube samples were briefly dipped twice in de-ionised water to remove any buffer salts and excess water was removed with a piece of filter paper. Holding at one end with a pair of tweezers, samples were plunge-frozen in liquid ethane and transferred to liquid-nitrogen cooled brass inserts. For fracturing, ethane-frozen samples were placed on a liquid nitrogen-cooled metal block and fractured with a cold razor blade before transferring to the brass insert.

The samples were freeze-dried overnight in a liquid nitrogen-cooled turbo freeze-drier (Quorum K775X), mounted on aluminium SEM stubs using conductive carbon sticky pads (Agar Scientific) and conductive silver paint, and coated with 35 nm gold using a Quorum K575X sputter coater. Examination of the collagen surfaces was carried out using a Verios 460 scanning electron microscope (FEI/ThermoFisher Scientific) run at an accelerating voltage of 2 keV and 25 pA probe current. Secondary electron images were acquired using either an Everhard-Thornley detector in field-free mode (low resolution) or a Through-Lens detector in full immersion mode (higher resolution).

SEM images were analysed using custom software to extract radius and orientation statistics for the collagen fibrils. The software is based on Frangi’s filter [57], a traditional filter for extracting tube-like features from images, with an approach to foreground segmentation that suppresses fibre bundles. Statistics were sampled from pixels in the skeleton of this (the centreline of the segmented fibrils). Radii, the filter scale at which the pixel responded most intensely, were extracted from images at 20,000x magnification, and orientations, the direction of the eigenvector associated with the near-zero eigenvalue of the smoothed Hessian, at 1,000x. Pores were extracted from images at 1,000x magnification by using the distribution of pixel intensities in the identified fibres to set a foreground threshold; combining this with the fibre foreground ensures that only dark regions are identified as pores, whilst helping to suppress background fibres. To ensure robustness to noise we follow the established approach [58] of morphological closing, but note that this introduces a minimum pore size. Furthermore, attempting to prevent pores from merging in some regions by setting the parameters controlling foreground segmentation to extreme values will completely close off actual pores. Hence, we set a minimum and maximum area by inspection and reject pores outside of this range. A further discussion of the image analysis can be found in the supporting information.

#### Second-harmonic generation imaging

Planar strips of collagen were cut from the full length of the tubes. Samples were mounted between two #1.5 glass coverslips, supported by a silicone spacer, and the coverslip ends were taped together. An inverted Leica TCS SP5 acousto-optic beam splitter (AOBS) multiphoton laser scanning microscope equipped with a Modelocked Ti:Sapphire Laser was used for second-harmonic generation (SHG) imaging, with the excitation laser (Ti:Sapphire) tuned to 880 nm. At this specific excitation wavelength, the wavelength of the emitted SHG signal was at 440 nm. A 20x objective was used to focus the excitation beam on the sample and to collect backscattered emission signals which were delivered through the AOBS and the fully opened confocal pinhole (600 μm) to the spectral detection system (descanned pathway). Highly specific SHG images were obtained through a 430-450 nm emission bandwidth channel detected on a hybrid detector. Tiled images were subsequently analysed using ImageJ.

#### Uniaxial tension

Tensile testing was carried out with an Instron desktop testing machine, fitted with a 5 N load cell. The tubes were cut along their length and ‘unwrapped’ so that they were planar. Using a metallic die, rectangular dogbone samples (*n* ≥ 3) were cut along the long axis of the unwrapped tubes, with gauge sections 7 mm long, 3 mm wide and 1.5 mm thick. For tensile testing of crosslinked samples, dogbone specimens were cut from the tubes prior to crosslinking. The samples were mounted into paper tabs with a central cut-out, using superglue. The top end of the paper tab was gripped using a bulldog clip and the bottom end using a specially designed vice clamp grip that used a pivot action to initially grip the sample and a clamping screw to tighten the vice. The jaws of the grip were rubber-faced to prevent sample slippage. Prior to testing, cuts were made from each side of the central cut-out. The crosshead displacement was measured using a linear variable differential transformer (LVDT). All tests were conducted in displacement control at a rate of 1 mm/min.

#### Burst pressures

The burst pressures of 10 mM genipin-crosslinked tubes (*n* ≥ 3) were measured, with a lumen diameter of 5 mm and wall thickness of 1.5 mm. An industrial pressure sensor (rated to 50 psi, Mouser Electronics) and a 5 mL syringe were interfaced to the collagen constructs via a pair of stainless steel tubes, supported by a pair of machined adapter pieces (supporting figure S5a). Heat shrink tubing (RS Components) was positioned onto the steel surface and the collagen tube placed over this. A further layer of heat shrink tubing was heated onto the ends of the tube to create a tight seal. A gap of 20 mm was maintained between the stainless steel tube interface at each end. A custom-made pressure measurement control system (supporting figure S5b) consisted of an instrumentation amplifier, Arduino process controller and serial data logger software (Termite). Pressure generated by a water-filled syringe was ramped up gradually until the tube burst, and the maximum pressure was recorded at the burst point.

#### Suture pullout forces

The forces during suture pullout were measured using an Instron desktop testing machine, fitted with a 5 N load cell. 10 mM genipin crosslinked tubes, with a lumen diameter of 5 mm and wall thickness of 1.5 mm, were cut open longitudinally to obtain 10 mm wide x 5 mm long sections (*n* = 6). The sections were clamped at the edge located opposite to the suture. A 7-0 polypropylene suture was threaded through the specimens (using a swaged needle) and both ends of the suture gripped into another clamp. The suture hole was centred with respect to the specimen width and its distance from the specimen end was 2 mm. The crosshead displacement was measured using a linear variable differential transformer (LVDT). Tests were conducted in displacement control at a rate of 3 mm/min until final failure, characterized by suture pullout.

#### Acid solubility

Representative samples (*n* ≥ 3), cut from genipin-crosslinked densified collagen tubes, were immersed in 100 mM acetic acid solution. A change in mass was measured over 7 days, using a mass balance with a precision of 0.1 mg.

### Cell culture and characterisation

#### Cell lines

Human liver cancer cells (HepG2s) were cultured in Dulbecco’s Modified Eagle Medium (DMEM) with 10 % FBS. Green fluorescent protein (GFP) and Red Fluorescent Protein (RFP) human umbilical vein endothelial cells (HUVECs, Promocell), normal human lung fibroblasts (NHLFs, Promocell) and mesenchymal stem cells (MSCs, Promocell) were cultured in their respective proprietary media (Promocell). Smooth muscle cells (SMCs) were derived from human embryonic stem cells (H9) via the neural crest or lateral plate mesoderm intermediates using established protocols [59, 60] and were subsequently cultured in standard medium (DMEM with 10 % FBS). Cells were cultured until 90 % confluency and detached from T75 flasks using TrypLE (Life Technologies), and subsequently centrifuged at 250 g for 5 min and counted using a hemocytometer.

#### Cell seeding into tube walls

Densification experiments where cells were encapsulated into the walls of the tubes were performed at 2-8 × 10^6^ cells per tube, using collagen tubes with luminal diameters of 2 and 5 mm. Tubes with cells encapsulated in the bulk were prepared using either HepG2s, NHLFs, GFP HUVECs and RFP HUVECs, or SMCs. The collagen precursor solution was prepared as above, subtracting a volume for the addition of cells later, and was subsequently centrifuged at 2000 g for 5 min. Approximately 1 mL of collagen gel precursor solution, without cells, was poured into the funnel to fill the cylindrical section. Doing so prevents the clogging of the membrane with cells and results in all the cells being present in the final tubular construct. The resuspended cell pellet was then pipetted on top of the precursor gel solution. Using a 1000 µL pipette tip, the cells were slowly mixed into the gel precursor solution, and the solution was then poured into the funnel assembly as described above and gelation and densification process continued as standard. For densified collagen tubes with two distinct cell domains, two 10 mL collagen precursor solutions were prepared. Prior to adding cells, 0.5 mL from each precursor solution was transferred into the chamber. GFP and RFP HUVECs were subsequently stirred into their respective precursor solutions and slowly poured into the chamber, one after the other.

#### Luminal surface seeding

HepG2s were used for experiments in which cells were seeded upon the luminal surface of the tubes. Using a two T75 flasks, 50 × 10^6^ HepG2 cells were pelleted and resuspended in 50 µL of DMEM with 10 % FBS. The 2 mm luminal diameter collagen tube was cut to length using small scissors and a sterile metal 2 mm diameter dowel pin (Accu) was positioned in the lumen at one end. Immersing the tube in culture medium, cells were transferred to the tube using a 1 mL syringe fitted with a 27 G × 1.5 in needle (Medisave). The cell-laden tube was left in a 37 °C incubator for 6 h and then a second volume of HepG2s (50 × 10^6^ cells) was added. Over several days of static culture, the attached HepG2s proliferated on the luminal surface to form a confluent layer.

For surface cell viability, an alamarBlue assay (Serotec, Oxford UK) was used as an indicator of metabolic function and proliferation. This assay contains a redox indicator; a blue, cell-permeable resazurin-based solution that is non-toxic and non-fluorescent. Upon entering living cells, resazurin is reduced to resorufin, a compound that is red in colour and fluorescent. SMCs were seeded on 1 and 10 mM genipin crosslinked collagen samples, cut into 4 mm diameter disks using a biopsy punch. PDMS and tissue culture plastic (TCP) of the same cross-sectional area served as the negative and positive controls respectively. SMCs were seeded at 2.6 × 10^4^ cells/cm^2^ in 5 µL droplets and incubated for 3 h to allow for cell attachment. After incubation, the samples were moved to 24-well plates and immersed in 1 mL of culture medium. The SMC surface-seeded samples were incubated for 4 h with fresh culture medium supplemented with 10 vol% alamarBlue reagent. Following incubation, 100 µL medium from each well was transferred to a 96-well microplate and replicated 3 times. Three independent experiments were conducted with four replicates from each group (*n* = 4). Fluorescence (excitation 530 nm, emission 590 nm) was measured on a Fluostar Optima microplate reader. The % reduction of alamarBlue was calculated using the following formula:

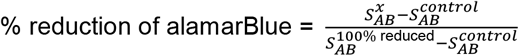

where 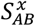 is the alamarBlue fluorescence signal at day x, 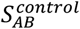 is the signal from the control, the culture medium supplemented with 10 vol.% alamarBlue reagent, and 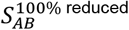 is the signal of the 100 % reduced form of alamarBlue obtained by autoclaving the control at 121 °C for 15 min.

#### Perfusion culture

Closed-loop perfusion of a cell-ladened collagen tube was undertaken using a custom-made apparatus. Densified collagen tubes were produced with an inner diameter of 2 mm and a wall thickness of 0.75 mm, using 2.5 mg/mL collagen gel (4 mL) containing 5 × 10^6^ SMCs. The densification process yielded a tube of final cell concentration of 2.5 × 10^7^ SMCs/mL embedded uniformly in the walls of the tube. A custom-made perfusion chamber was machined, and the tube mounted across a pair of stainless-steel hypodermic tubes, using microfluidic ferrules to create a tight seal. Prolene suture was used to create a seal between the hypodermic inlet and collagen tube. Filling the chamber with cell medium, the lid was tightly fixed in place. A peristaltic pump and cell medium reservoir (40 mL) were used to undertake closed loop perfusion at 600 mL/hr for 10 days, changing the cell medium perfusate on alternate days. An estimate of cell number (and associated cell density) at the end of the 10-day experiment was undertaken through manual cell counting (ImageJ) using 5 µm thick cross-sectional slices which had been histologically stained with H&E.

#### Staining and imaging

Cell characterisation was performed via calcein AM staining (Life Technologies). For tube wall cell seeding experiments, cells were either stained prior to encapsulation or after the densification process. Briefly, calcein AM was resuspended at 4 mM in PBS. Cells plated in T75 flasks, or cell-seeded tubes, were washed in PBS and incubated in a 4 µM calcein AM solution in PBS, at 37 °C for 20 min. For luminal surface seeding experiments, HepG2s were stained prior to seeding. Tubes that had undergone perfusion culture were fixed in 4 % PFA for 24 h and subsequently dehydrated in ethanol and xylene, embedded in paraffin wax and sectioned into slices of 5 and 20 µm thicknesses. Histological (Masson’s Trichrome, H&E) and immunohistochemical stains (αSMA via fluorescent immunodetection, TAGLN via horseradish peroxidase (HRP)-mediated chromogenic immunodetection) were produced to illuminate the cell and collagen structures within the construct. Paraffin-embedded tissue sections were rehydrated and a heat-mediated antigen retrieval step was performed using a citrate buffer (pH 6.0) containing 0.05 % Tween 20. Tissue sections were subsequently permeabilized in a 0.25 % solution of Triton X-100 for 10 min. For chromogenic staining, an incubation with BLOXALL (Vector Laboratories) was performed at this stage. All sections were then blocked using 10 % goat serum in Tris-buffered saline (TBS) for 1 h. Next, primary antibodies (αSMA, 1:100, Bio-Techne; TAGLN, 1:100, Abcam) were added in TBS, containing 10 % goat serum, and incubated overnight at 4 °C. The following day, the samples were rinsed three times using TBS. For fluorescent immunostaining, an Alexa Fluor 568-conjugated secondary antibody was added (1:1000, Abcam) in TBS containing 10 % goat serum, for 1 h. Unbound secondary antibodies were subsequently removed by washing samples three times in TBS. To counter collagen autofluorescence, the samples were incubated in 70 % ethanol containing Sudan Black B (0.1 %, Fisher Scientific) and subsequently washed with 70 % ethanol and then PBS. A nuclear counterstain was conducted using Hoechst 33342 (1:2000, Life Technologies) in PBS for 5 min, followed by washing three times using TBS. For chromogenic immunostaining, a HRP-conjugated secondary antibody was added (1:1000, Sigma) in TBS containing 10 % goat serum, for 1 h. Unbound secondary antibodies were subsequently removed by washing samples three times in TBS. Samples were next incubated in ImmPact DAB Substrate, Peroxidase (HRP, Vector Laboratories) for 8 min and washed with water, and then counterstained using Hematoxylin (Vector Laboratories). Samples were mounted using Vectamount AQ Mounting Medium (Vector Laboratories). Samples and sections were imaged using a bright-field and an epi-fluorescence microscope (Zeiss Axio-Observer.Z1).

### Luminal surface patterning process

Inverse 3D printed wax models (Solidscape) were prepared using a standard method [61]. 20 %(w/v) high bloom, porcine gelatin solution was poured into the wax moulds and degassed using a vacuum pump. A polished nylon rod, 2 mm in diameter with a cutaway section at one end with 0.5 mm diameter, was then fitted through the wax mould and the assembly left at 4 °C to allow for gelation of the gelatin. The construct was subsequently immersed in acetone solution to dissolve the 3D printed wax mould and washed in PBS. The standard collagen gelation and densification process were subsequently undertaken. Following densification of the collagen gel, the tube with patterned gelatin was placed in a PBS bath at 37 °C to melt the gelatin and allow it to diffuse away, leaving a patterned collagen tube.

### Statistical analyses

In all figures, data are expressed as the arithmetic mean ± standard deviation over *n* independent replicates, detailed in figure captions. Individual data points are plotted where appropriate. AlamarBlue data analysis was performed using IBM SPSS Statistics 28. A Levene’s test showed that the variances were not homogenous so data were analysed for statistical significance using the Kruskal-Wallis test and the Games-Howell post hoc test. Differences were considered statistically significant at *p* < 0.05. n.s. means no statistically significant differences were measured.

## Author Contributions

AWJ and AEM designed research; AWJ conceived of, designed, and performed the densification method, analysed and interpreted associated results; AWJ, FC, AAG: architectural characterization ; AWJ, SRE, SB, AST: mechanical testing; AWJ, FC, HD, JO, AGJ: cell experiments. AWJ and AEM data interpretation and figure preparation; AWJ wrote the main manuscript with AEM; KSP, SS, AEM: supervised the work. All authors: critical revision and final approval of the manuscript.

## Acknowledgements

Financial support for this work has come from the Engineering and Physical Sciences Research Council (EPSRC, EP/R511675/1), the Isaac Newton Trust, the Rosetrees Trust (M787) and the Wellcome Trust Institutional Translational Partnership Award 222062/Z/20/Z. Financial support for JO has been provided via a WD Armstrong Studentship. Support for SS and HD via BHF awards (FS/18/46/33663 and RG/17/5/32936) is gratefully acknowledged. We wish to acknowledge Fadwa Joud of the Cancer Research UK, Cambridge Institute for second harmonic generation microscopy, Peter J. Holt of the Wellcome–Medical Research Council Cambridge Stem Cell Institute, for smooth muscle cell culture, Stefan Savage of the Department of Engineering for machining the Perspex funnels, Peter W. Justin for design and manufacture of the burst pressure measurement apparatus, Karin Mueller of the Cambridge Advanced Imaging Centre (CAIC), for SEM processing and imaging, and Louise Howard of the Department of Pathology, for histological processing and staining. The authors are grateful to Prof. Michael Sutcliffe for discussions on suture testing.

## Data availability

The datasets generated and analysed during the current study are available to download from Mendeley Data, V1: https://doi.org/10.17632/3r5knpm6g5.1.

