## Supporting Information for "Densified Collagen Tubular Grafts for Human Tissue Replacement and Disease Modelling Applications"

### SUPPPORTING INFORMATION

### SEM IMAGE ANALYSIS

After initial attempts at using standard Fiji [1] plugins Frangi and Tubeness, and the more specialised DiameterJ [2], as well as other options for analysing tube-like data (AngioTool [3]) failed, a custom toolkit was written to analyse the images. Some reasons for failure include:

- Fibres falling below the minimum diameter for some plugins, and the foreground containing merged fibre bundles.
- Many fibres merging together and giving responses at higher diameters.
- Edge highlights giving a response at lower diameters.

This was leading to poor skeletonisation and clearly incorrect results for diameter and connectivity.

#### Fibres

As the images obtained are cluttered and noisy with relatively low fibre diameters for the existing software to operate robustly, as well as fibre bundles which we wish to ignore, the framework focuses on suppressing spurious signals by aggressively rejecting candidate foreground (fibre) pixels. Due to the variance in pixel depths, extracting fibre connectivity data from these 2D images is infeasible as this would require identifying overpasses/connections correctly. From the foreground fragments which are accepted, a skeleton is created, and statistics are sampled at these points, giving an estimate of the fraction of fibres in the image at each diameter and orientation.

The processing pipeline is based on Frangi's filter [4], a classical filter that looks at the eigenvalues of the Hessian matrix of the image, smoothed at a given length scale. The length scale is the fibre radius, as the second derivative of the Gaussian has its zero-crossings at this point. Orientation is extracted from the eigenvector of the Hessian associated with the near-zero eigenvalue, modulo pi (as we do not care about the sign of the direction vector). The traditional Frangi filter is the maximum intensity projection (MIP) over several length scales – in this work we increment by 1 px (pixel), assigning fibre diameter in 2 px intervals. All basic components used in the processing pipeline (filters, thresholding methods, morphological operations) are from the scikit-image package [5].

To sparsely sample fibre pixels, we attempt to imitate how a human may select the most prominent fibres and reject the merged bundles. We start by inspecting several statistics at each filter length scale:

- *Response*: the sum over the output image at that scale (Frangi offered the interpretation of his filter as the probability of a pixel belonging to a vessel-like structure of given radius, in which case this corresponds to the average probability of a pixel in the image being a fibre at that scale).
- *Dominance*: the number of pixels in the MIP over all scales that were taken from this scale.
- *Contribution*: the sum over those pixels (the total "fibre probability" contributed at this scale).
- *Intensity* = *Contribution / Dominance*: the average contribution per pixel.

Some example images at low and high magnification with their Frangi MIPs are shown in figures S1a,b and S2a,b, and the associated filter statistics in figures S1g and S2g. We see that there are many pixels corresponding to bundles and other large-diameter structures in the output image, leading to high dominance and contribution at the largest filter length, but the intensity is focused at the lower length scales which agree with manually measured samples. We therefore generate a foreground threshold using the MIP of filtered images at length scales above a given intensity threshold (default 90 % of maximum), using Otsu's method [6] (setting the threshold to 0 recreates Otsu thresholding the traditional Frangi MIP) – see figure S1c and S2c. Spurious vessels (e.g. those detected in the edge effects of larger bundles) are then removed using morphological opening – there is a trade-off here between rejecting noise and accepting fibres, and this was set by inspection to a 2 px opening radius (figure S1d and S2d). The accepted fibre fragments can be seen overlaid on the original images in figure S1e and S2e with the skeletons shown in figure S1f and S2f.

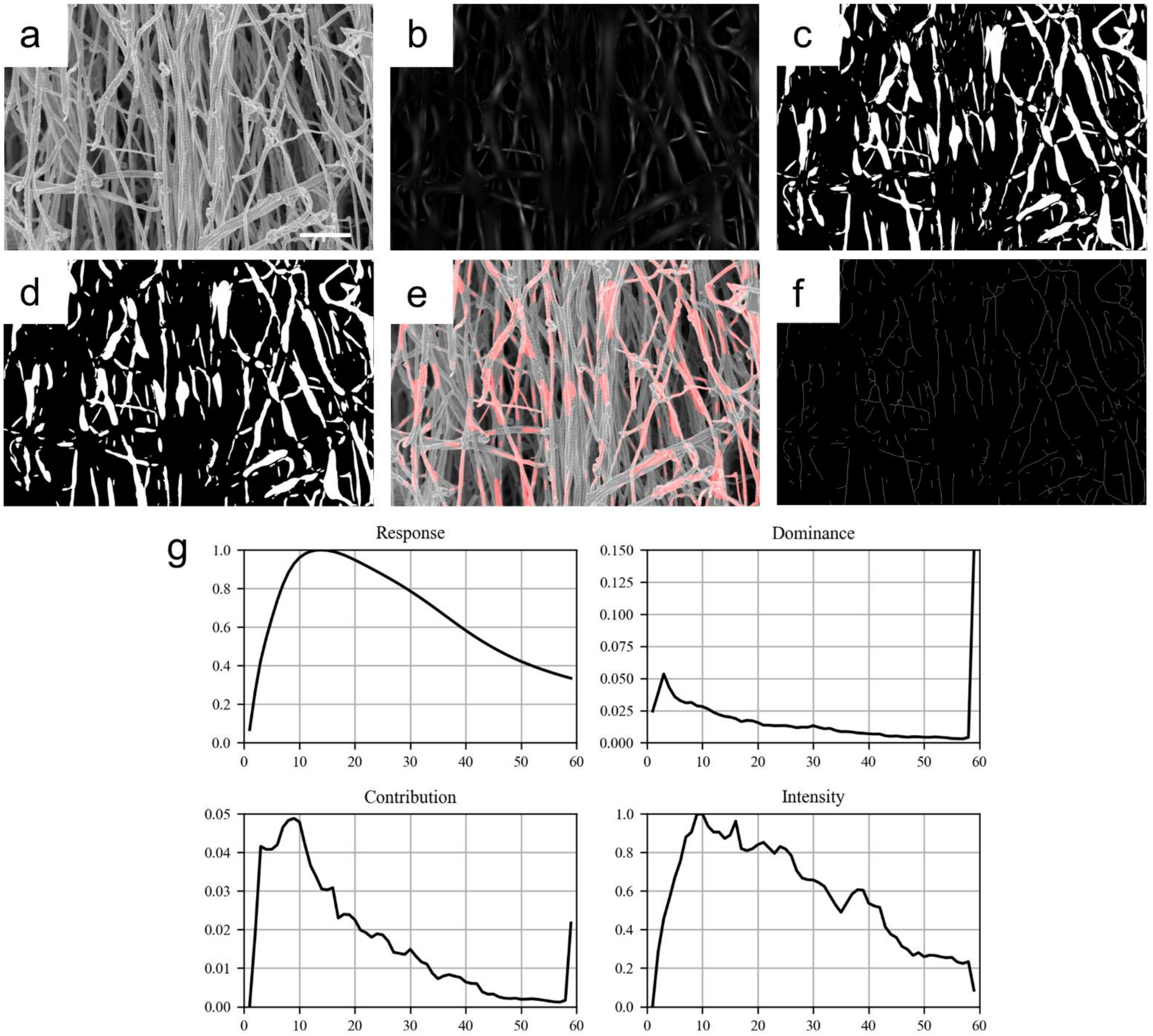


**Figure S1.** Low magnification SEM image analysis. **(a)** Low magnification fibres. Scale bar: 10 µm. **(b)** Frangi filter MIP over length scales 1-60 px. **(c)** Intensity-based thresholding foreground. **(d)** Morphological opening of the foreground to remove spurious fibres. **(e)** Accepted “fibre-fragment” foreground overlaid on original image. **(f)** Fibre skeleton, from which statistics were sampled. **(g)** Frangi filter statistics over length scales 1-60 px, again showing the peak at the largest scale but without a spurious peak at the lowest scale due to the low magnification – fibres are only a few pixels wide at this magnification.

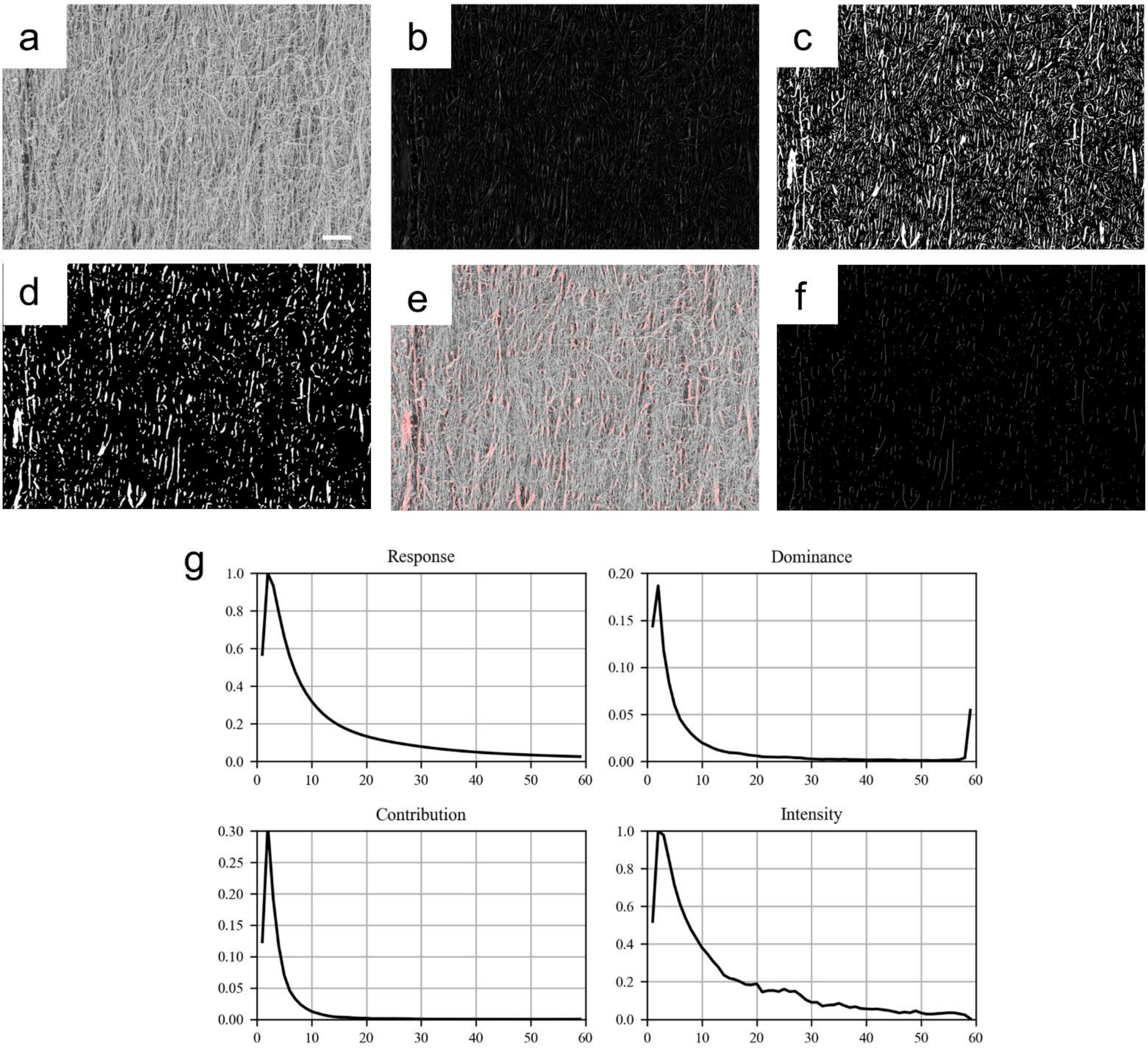


**Figure S2.** High magnification SEM image analysis. **(a)** High magnification fibres. Scale bar: 1 µm. **(b)** Frangi filter MIP over length scales 1-60 px. **(c)** Intensity-based thresholding foreground. **(d)** Morphological opening of the foreground to remove spurious fibres. **(e)** Accepted “fibre-fragment” foreground overlaid on original image. **(f)** Fibre skeleton, from which statistics were sampled. **(g)** Frangi filter statistics over length scales 1-60 px, showing high response from bundles at the highest length scale and edge highlights at the lowest scales; these are suppressed when considering intensity.

<u>Pores</u>

For pore analysis of cross-sectional slices, we use a similar approach. However, we now wish to reject pores, meaning that we wish to oversample the foreground. Further, regions of high, constant intensity are not valid as pores, therefore we cannot simply rely on the Frangi filter outputs.

The approach used is to start from the same foreground segmentation of fibres (figure S3d,e), but this time morphologically closing instead of opening (figure S3f). To reject constant high-intensity foreground elements, the intensity of pixels in this fibre foreground are sampled and the threshold is set as $T = mean(I) - k\ std(I)$, where $k$

defaults to 1 and aims to ensure that enough of the foreground pixels are recognised – the idea is that these "intense fibres" correspond to top-layer fibres, a subset of the solid mass in the top layer, and it is the gaps between the solid mass of this layer that we wish to measure (figure S3a-c). This value is then used to segment the original image, morphologically closed, and the two foregrounds are merged (figure S3g-i). Note that we have again a trade-off:

- Closing removes small pores, and there is a minimum pore size after this is complete (the stencil used in the erosion applied to a single pixel). We therefore remove pores below a certain size.
- In regions of noise or low signal, pores can merge. Countering this (lower threshold, higher closing radius) eliminates the genuine pores at smaller scales.

Parameter values were therefore determined by manually measuring a number of pore diameters, then tweaked based on visual inspection of the output. The resulting foreground can be seen in figure S3b,c. The pores were then labelled, and data (area, orientation, lengths) returned by the *regionprops* function.

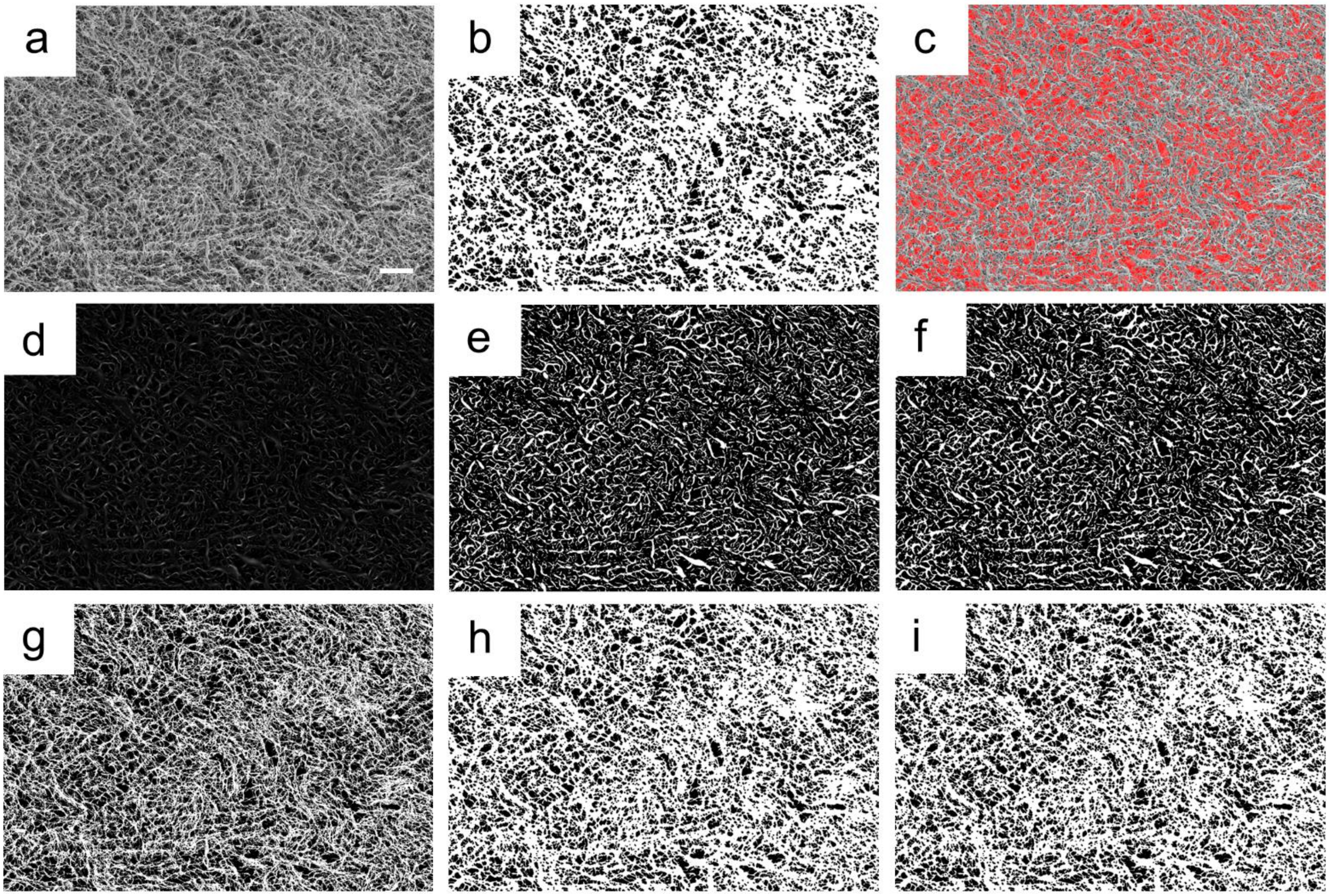


**Figure S3.** Pore size image analysis. **(a)** Cross sectional image of fibres with pores. Scale bar: 10 µm. **(b)** The identified foreground, with small holes removed. **(c)** Foreground inverse overlaid on original image, highlighting pores. **(d)** Frangi filter MIP over length scales 1-60 px. **(e)** Intensity-based fibre foreground. **(f)** Fibre foreground, morphologically closed. **(g)** Original image foreground, thresholded using distribution of intensities in *top-layer* fibres identified in previous step. **(h)** Morphological closing of original image foreground. **(i)** Merged foreground images, before removing holes below the minimum identifiable feature size.

**Second Harmonic Imaging of Sloped Funnel Collagen Tubes**

Second harmonic imaging was undertaken to observe fibre orientation and density. Uncrosslinked collagen tubes were cut open longitudinally and the outside surface of the collagen was imaged. The output grayscale values were extracted from the second harmonic imaging data, which represented the density of the collagen fibrils. The use of the sloped funnel moulds shows a density gradient of fibrils along the length of the tube (figure S4). The maximum collagen density was measured at the top end of the collagen tube, where the sloped funnel was widest. Drops in signal along the length were associated with difficulties in keeping the sample completely flat between the coverslips, and with minor surface defects associated with bubble formation during tube manufacture.

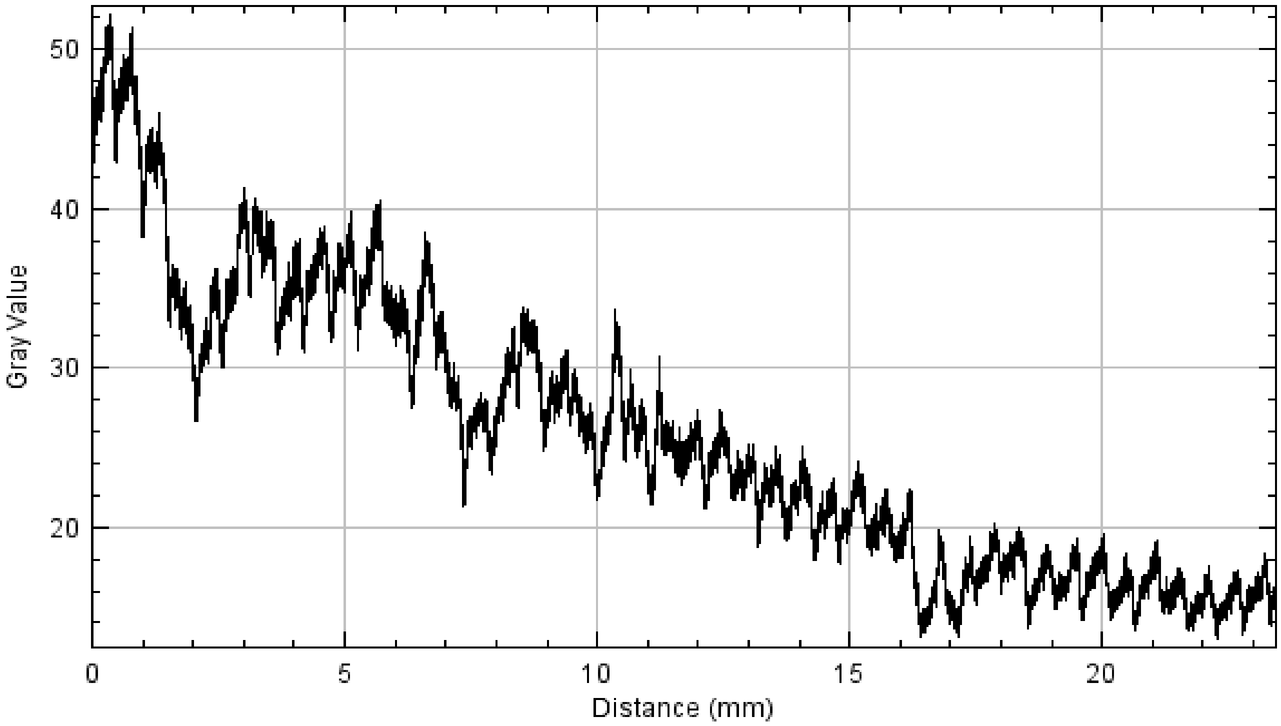


**Figure S4:** Variation of collagen density along tube length.

**Burst Pressure Testing**

Additional information on burst pressure tests conducted on densified collagen tubes.

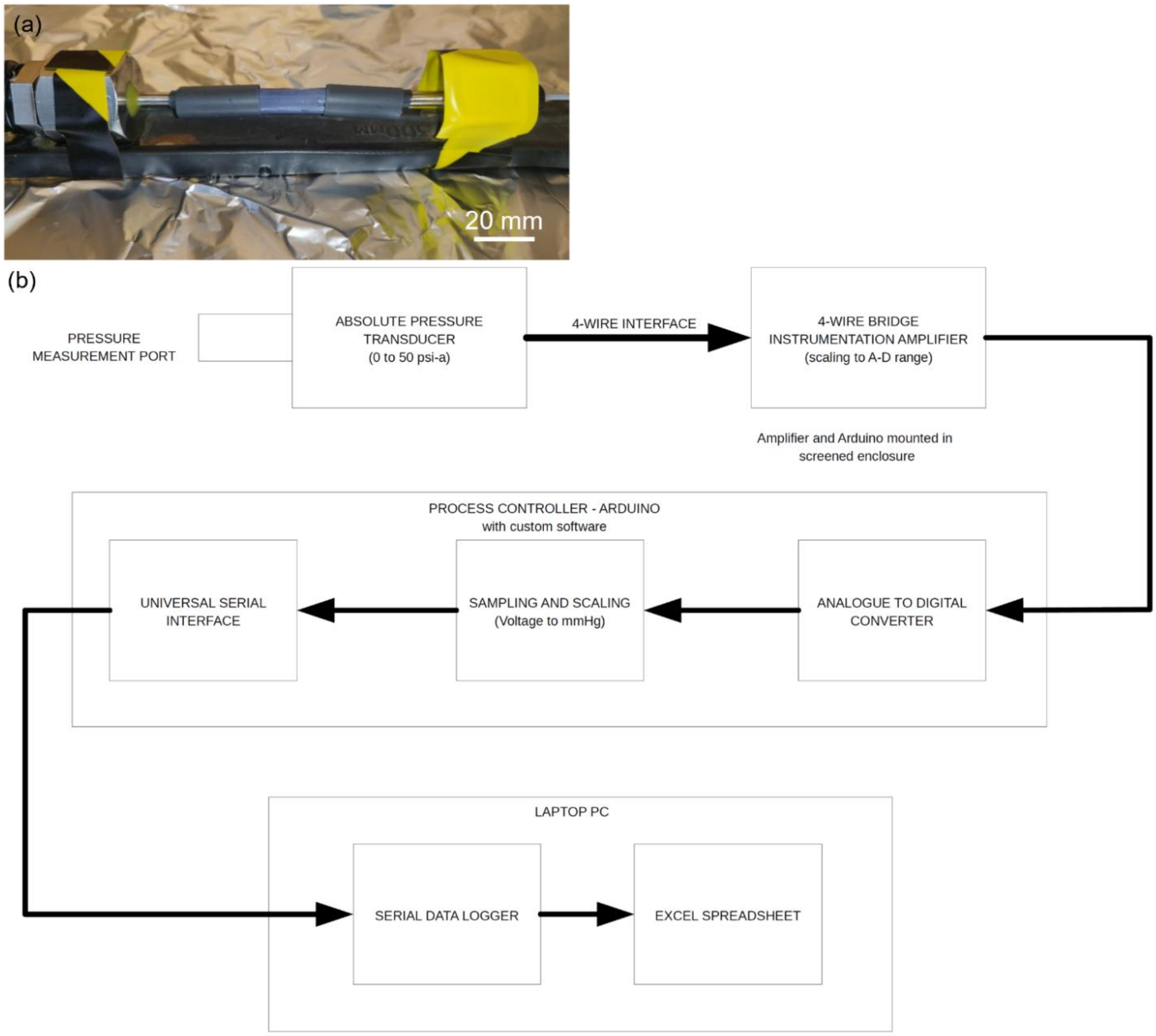


**Figure S5: (a)** Collagen tube mounted on a pair of stainless steel tubes which were connected to the pressure sensor and syringe at each end by a pair of machined adapter pieces. Heat shrink tubing was used to seal the ends. **(b)** Pressure measurement system.
